# Statistical structure of the trial-to-trial timing variability in synfire chains

**DOI:** 10.1101/2020.03.21.001503

**Authors:** Dina Obeid, Jacob A. Zavatone-Veth, Cengiz Pehlevan

## Abstract

Timing and its variability are crucial for behavior. Consequently, neural circuits that take part in the control of timing and in the measurement of temporal intervals have been the subject of much research. Here, we provide an analytical and computational account of the temporal variability in what is perhaps the most basic model of a timing circuit, the synfire chain. First, we study the statistical structure of trial-to-trial timing variability in a reduced but analytically tractable model: a chain of single integrate-and-fire neurons. We show that this circuit’s variability is well-described by a generative model consisting of local, global, and jitter components. We relate each of these components to distinct neural mechanisms in the model. Next, we establish in simulations that these results carry over to a noisy homogeneous synfire chain. Finally, motivated by the fact that a synfire chain is thought to underlie the circuit that takes part in the control and timing of zebra finch song, we present simulations of a biologically realistic synfire chain model of the zebra finch timekeeping circuit. We find the structure of trial-to-trial timing variability to be consistent with our previous findings, and to agree with experimental observations of the song’s temporal variability. Our study therefore provides a possible neuronal account of behavioral variability in zebra finches.

## I. INTRODUCTION

Timing is critical for many behaviors, such as speech production, playing a musical instrument, or dancing. However, even the most stereotyped animal behaviors are significantly variable from one iteration to the next. This so-called trial-to-trial variability is ubiquitous and may serve important functions in motor learning and adaptation [1–3]. Its sources have therefore been of great interest to neuroscientists [2, 4].

As behavior is controlled by the nervous system, it is natural to look for the source of some of this variability in the variable activity of neural circuits involved in the production of behavior [2, 4–7]. Indeed, the neural mechanisms underlying behavioral timing have been extensively studied experimentally [2, 8–12], establishing links between temporal variations of behavior and that of neural activity [13–20]. This neural variability could result from multiple sources such as stochastic events at the level of ion channels [21], synapses [22], and neurons [23]; chaotic activity of neural networks [24, 25]; and sensory inference errors [5]. Therefore, understanding the mechanisms and structure of timing variability in neural circuits is necessary for understanding variability in behavior.

In this paper, we study temporal variability in one of the most basic neural network models of timing, the synfire chain [26–28]. The synfire chain is a feedforward network composed of pools of neurons that produce traveling waves of synchronized spiking activity, as illustrated in Figure 1. The synchronization of spikes within a pool, and the sequential propagation of activity across pools (Figure 1b) allow the synfire chain to serve multiple timing functions in a very natural way. First, it can be used to keep time by simply counting the pool which the spiking activity has reached. Second, it can be used to produce precisely timed intervals defined by the time elapsed between when activity arrives at a given pool and when it arrives at a subsequent pool (Figure 1b). The synfire chain can sustain activity indefinitely given sufficiently many pools [27, 29], or by arranging it in a circular topology such that the final pool connects back to the first one [29]. While the synfire chain is theoretically well-studied [27, 29–31], and experiments support its existence in biological systems [32], a theoretical account of its temporal variability is still lacking.

**FIG. 1.**
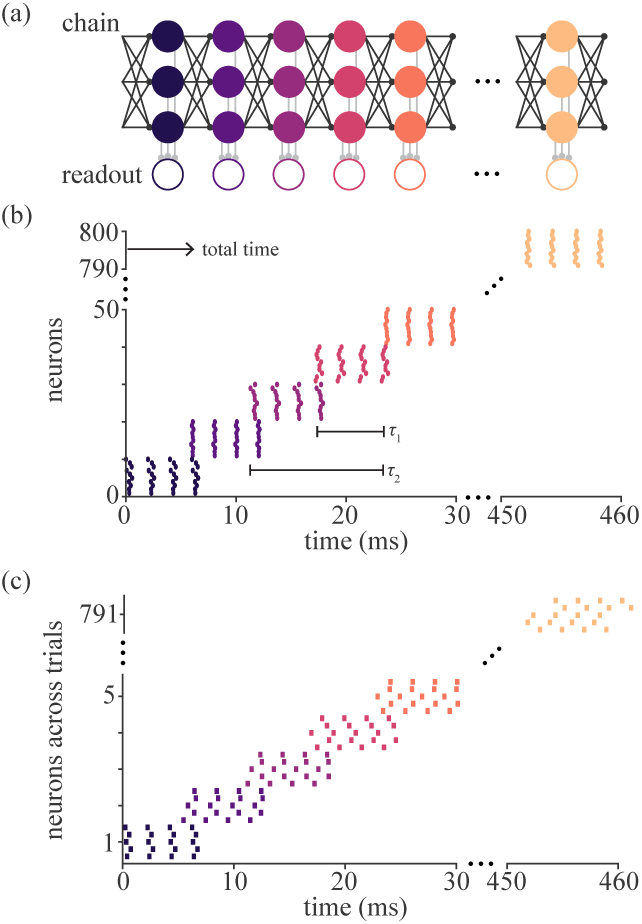
Timekeeping and trial-to-trial variability in synfire chains. (a) The synfire chain is a feedforward network of pools of neurons. In the figure, circles represent neurons and arrows represent synapses. A simple scheme to measure time intervals composed of *K* successive pools, is to mark the beginning and end of the interval by the first spike of readout neurons corresponding to first and last pools. (b) Spike trains produced by the synfire chain show a synchronized activity that progresses pool-by-pool. Each point corresponds to the time of a spike produced by a neuron in the chain. Color denotes membership in a layer, each of which consists of a pool of 10 neurons. Various time intervals are shown. (c) Trial-to-trial variability in the synfire chain. Spike times of six neurons from different pools are plotted in different colors. Five different trials for each neuron are shown.

We are interested in the trial-to-trial variability in the timing of neural activity of a synfire chain. Such variability arises from the millisecond-scale tempo differences across multiple propagations of the spiking activity in the chain. We will focus on trial-to-trial variability caused by the inherent noise and fatigue in the neural system.

While it might seem small, millisecond-scale neural variability has been experimentally shown to correlate with behavioral variability at the same timescale in songbirds [16, 18, 20]. This finding is especially relevant since experiments support the existence of a synfire chain architecture in the songbird premotor cortex [32] for millisecond-scale precise time-keeping of the birdsong, with total song durations of few hundreds of milliseconds [13, 33]. Therefore, our findings may have direct implications for behavioral timing. Indeed, we will show that the statistical structure of temporal variability in a synfire chain can possibly explain some of the salient features of temporal variability in birdsong [33–35].

We address these questions first in a simplified and analytically solvable model of trial-to-trial variability in a chain of individual neurons (Section II). We derive analytical expressions that describe the magnitude and statistical structure of temporal variability in terms of inherent neural noise and fatigue and verify our results with simulations. We use a generative model introduced by Glaze and Troyer [35] to decompose the variability covariance matrix into three components: independent, global and jitter. Next, we address temporal variability in a noisy homogeneous synfire chain, which includes multiple identical neurons per pool, by numerical simulations (Section III). We show that our results from the analytically tractable model qualitatively carry over to this more complex model. We study the dependence of the various components of variability on the number of neurons per pool of the chain. Further, we relate the distinct neural mechanisms in the model to the different components of variability obtained from the generative model. Finally, we provide an application of our results to birdsong. In zebra finches, experimental studies support the existence of a synfire chain structure in the premotor nucleus HVC [32], which takes part in the production and timing of the birdsong. We simulate a biologically realistic synfire chain model [32] and show that the statistical structure and magnitude of its variability is consistent with that observed in the analytically tractable and homogeneous synfire chain model, and the zebra finch song (Section IV).

## II. TRIAL-TO-TRIAL TIMING VARIABILITY IN A CHAIN OF SINGLE NEURONS

In this section, we describe the statistical structure of trial-to-trial timing variability in an analytically tractable model: a chain of *N* single leaky integrate-and-fire (IF) neurons. In this simple model, each neuron is driven by excitatory synaptic input from the previous neuron in the chain at time *t*_*ps*_, which we model by *I*_*s*_Θ(*t − t*_*ps*_), where Θ(*t*) is the Heaviside step function. Though it would be more biologically realistic to use a input kernel of finite duration, we make this analytically convenient choice as we are only concerned with first-spike times. We model the drive to each neuron from outside the chain by the sum of a constant current *I*_0_, and the noise due to synaptic transmission and other cellular processes by a zero-mean Gaussian process 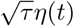 with autocorrelation *σ*^2^*τδ*(*t* − *t*′), where *τ* is the membrane time constant and *σ* controls the standard deviation of the noise [36, 37].

The sub-threshold dynamics of the membrane potential *V* of a given neuron in the chain is then governed by the Langevin equation [36, 37]

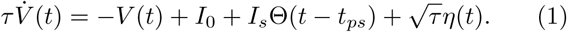

When the neuron’s membrane potential reaches a firing threshold *V*_*th*_, the neuron produces a spike and resets its membrane potential to *V*_*r*_.

### A. Local variability in a chain of single neurons

We want to study the variability in the first-spike times of successive neurons in the chain. This problem differs from the standard treatment of noisy and leaky IF neurons [36, 37] in that we are interested in trial-to-trial variability of intervals between different neurons’ spikes rather than long-time statistics of the intervals between spikes generated by a single neuron. However, we can map this problem to previous results in the literature [36, 37] by dividing it into two threshold-crossing problems. First, we can determine the probability distribution of a given neuron’s membrane potential at time *t*_*ps*_, using the fact that it receives no synaptic input before the previous neuron’s first spike. Then, given that its membrane potential at time *t*_*ps*_ is *V*_0_ with probability *P* (*V*_0_), we can think of the trial-to-trial variability in that neuron’s time to first-spike after *t*_*ps*_ as the variability in the inter-spike intervals of a single leaky IF neuron with *V*_*r*_ = *V*_0_.

To proceed we assume that *t*_*ps*_ is long enough such that *V* (*t*_*ps*_) has equilibrated. This assumption simplifies our calculations and will be validated when our final results are compared to simulations. Then, the solution to the first problem is given by the stationary distribution of the membrane potential, which was found in [36] to be

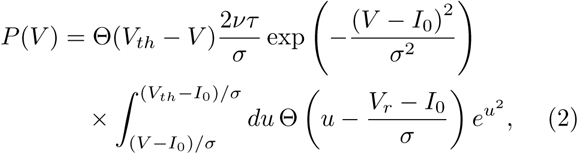

Where the firing rate *v* is given by

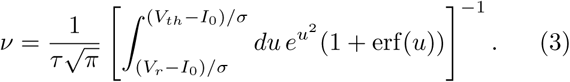

We will be interested in the limit of very low firing rates. This limit is relevant to propagation of spiking activity in synfire chains because a neuron in the chain spikes only when the spiking activity reaches the pool to which the neuron belongs and very rarely otherwise. Because noise is the main driver of firing in the absence of external input, the very low firing rate limit is given by assuming *V*_*th*_ − *I*_0_ ≫ *σ* [37, 38], which leads to the standard approximations (see Appendix A 1 and [36]) of the firing rate as

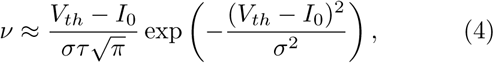

and the membrane potential distribution as

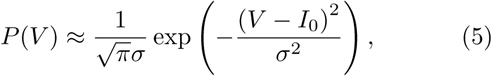

which is the stationary limit of an Ornstein-Uhlenbeck process with boundaries set at infinity.

We can calculate the mean and variance of the first-spike-interval, *T*_*fs*_, defined as the time elapsed from *t*_*ps*_ to the arrival of the first spike, using the mapping between our problem and that of determining the statistics of the inter-spike intervals of a single leaky IF neuron. Conditioned on *V* (*t*_*ps*_) = *V*_0_, these statistics are given by the standard expressions [36, 37]

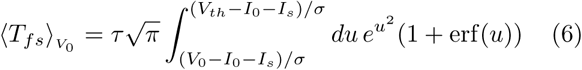

and

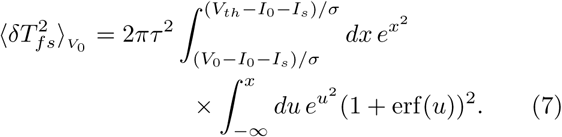

We can then combine these results to compute the mean and variance of the first-spike-interval across trials.

If we approximate the distribution of initial membrane potentials by the stationary Ornstein-Uhlenbeck process limit (5) in the low firing rate regime *V*_*th*_ − *I*_0_ ≫ *σ*, we obtain the lowest-order asymptotic expansions

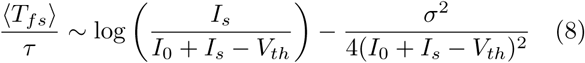

and

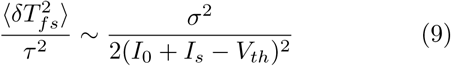

in the limit of large synaptic input *I*_*s*_ + *I*_0_ − *V*_*th*_ ≫ *σ* (a detailed derivation of these expressions is given in Appendix A 2). The scaling of this variability with *I*_*s*_ and *σ* for fixed *I*_0_ and *V*_*th*_ is illustrated in Figure 2 (see Section II D). In Appendix A 3, we also derive asymptotics for ⟨*T*_*fs*_⟩ and 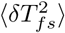 using the alternative approximation *P* (*V*_0_) ≈ *δ*(*V*_0_ − *I*_0_).

**FIG. 2.**
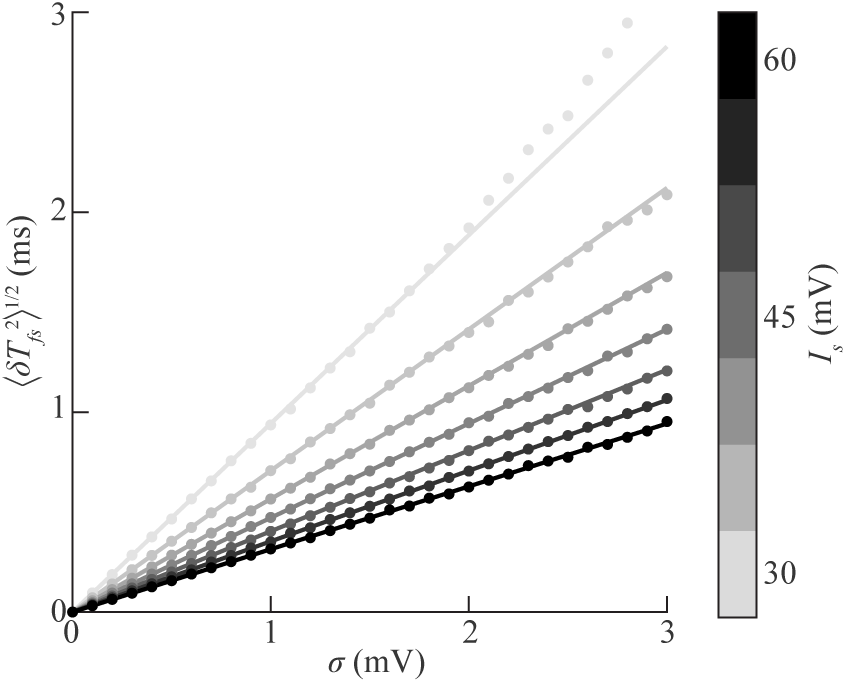
Scaling of local trial-to-trial timing variability in a chain of single neurons. The results of numerical experiment (Section II D) are shown as dots, and the solid lines show the asymptotic approximations obtained in Section II A. The ordinate shows the standard deviation 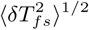 of the first-spike-interval, and the abscissa shows the standard deviation *σ* of the noise. Increasing values of *I*_*s*_ are indicated by darker shades of gray.

### B. Correlated variability in a chain of single neurons

Thus far, we have only considered sources of variability that are independent across neurons. However, in biological neural networks, there are many possible mechanisms that could introduce correlated variability, such as correlated external input [4, 7, 37, 38]. Here, we consider a simple model for correlated variability due to neural fatigue, i.e. a loss in a neuron’s excitability due to effects like synaptic depression [39].

In our model, the spiking threshold in a given trial is increased by some multiple *m* ∈ {0, 1, …, *m*_max_} of a small increment *δV*_*th*_ relative to the baseline threshold *V*_*th*_. We assume that, across trials, ⟨*m*⟩ is drawn from some distribution with mean *m* and variance ⟨*δm*^2^⟩. Working in the regime in which *m*_max_ *δV*_*th*_ ≪ *I*_0_+*I*_*s*_ − *V*_*th*_, relevant to neurons in a synfire chain which receive a barrage of inputs from the synchronous firing of presynaptic pool of neurons, we can use our previously-obtained asymptotic expansions (8, 9) of the mean and variance of the first-spike-interval conditioned on *V*_*th*_ to obtain

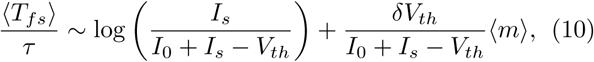

and

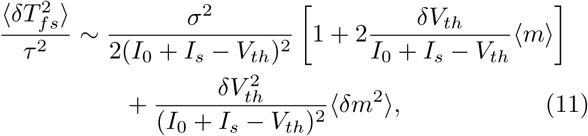

to lowest order in both *δV*_*th*_*/*(*I*_0_+*I*_*s*_ − *V*_*th*_) and *σ/*(*I*_0_+*I*_*s*_ − *V*_*th*_) (see Appendix A 2 for details). If we now consider two different neurons *a* and *b*, the trial-to-trial covariance of their first-spike-intervals 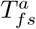 and 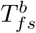 will be

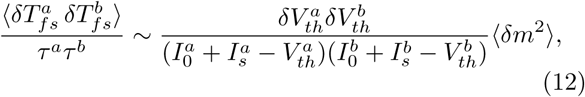

to lowest order in 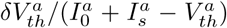 and 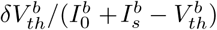.

Therefore, we obtain a model in which the trial-to-trial covariance matrix of the first-spike-intervals of neurons in the chain is the sum of a diagonal, local-to-a-neuron component and a rank-one global component. Concretely, if we assume for simplicity that all neurons in the chain are identical, the covariance matrix of the first-spike-intervals of the neurons in the chain is given as 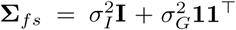, where 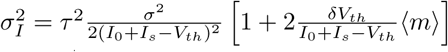 is the local component of the first-spike-interval variance, 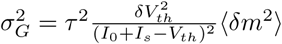 is the global component of the first-spike-interval variance, **I** is the identity matrix, and **1** is the ones vector. If the neurons were non-identical, *σ*_*I*_ and *σ*_*G*_ would no longer be scalar constants, and the covariance matrix would have the form 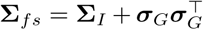 for a diagonal matrix **Σ**_*I*_ and a vector ***σ***_*G*_. This decomposition is illustrated in Figure 3a.

**FIG. 3.**
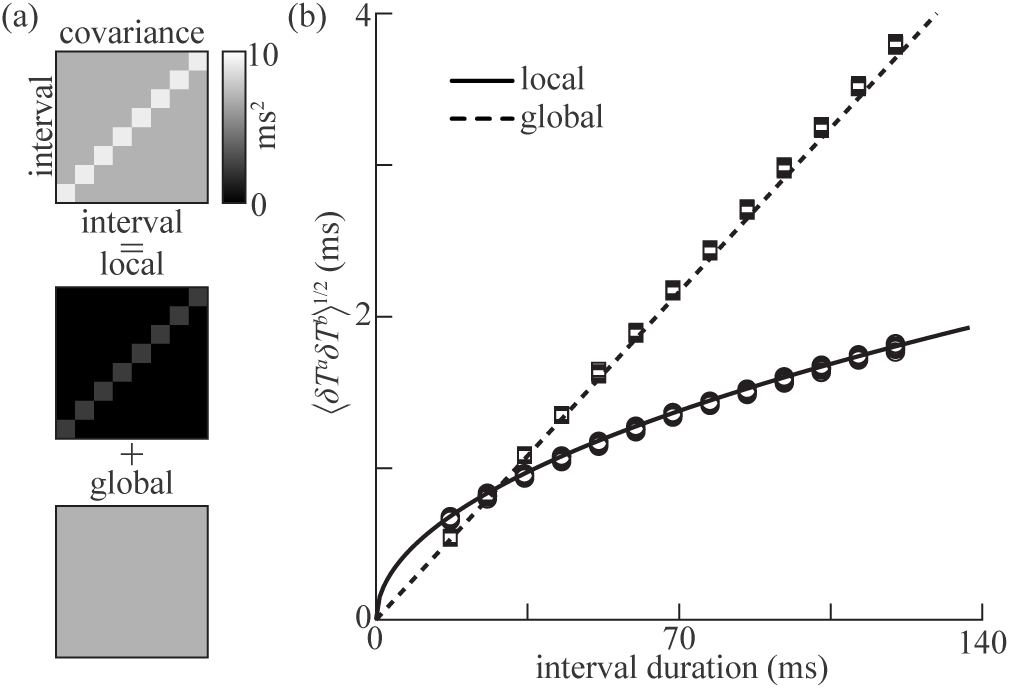
Correlated trial-to-trial timing variability in a chain of single neurons. (a) Schematic representation of the decomposition of the interval-interval covariance matrix into local and global components as described in Section II B (see expressions for 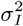 and 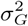). (b) Scaling of the local and global variability with interval duration. The results of numerical experiment (see Section II D) are shown by circles and squares for local and global variability, respectively, while the predictions of asymptotic theory (see Sections II B and II C) are shown by solid and dashed lines. Realizations of the random sampling used to generate intervals are plotted as individual markers.

In the above calculation, we assumed that one can perfectly read out the first-spike times from the neurons of the chain. However, it is unlikely that noise-free readout is possible in biological timekeeping systems. In the presence of readout noise, the first-spike interval covariance matrix will have an additional component that increases the variance of individual intervals and introduces negative covariance between adjacent intervals, a phenomenon known as timing jitter [35, 40]. Assuming for simplicity that the readout noise is homogeneous, additive, and independent of other forms of variability and has standard deviation *σ*_*J*_, the overall covariance matrix is given as

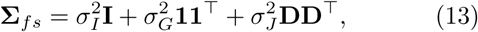

where **D** is the *N* × (*N* − 1) bidiagonal differencing matrix with ones along the diagonal and negative ones along the subdiagonal: *D*_*jk*_ = *δ*_*jk*_ − *δ*_*j*(*k*−1)_.

### C. From mechanism to statistical models of timing variability

In the preceding sections, we have characterized the timing variability inherent in a simple neural model. This calculation showed that the covariance matrix of the intervals between the first spikes of successive neurons in the chain could be decomposed into a diagonal local component, a rank-one global component, and a structured component resulting from imperfect readout of spike times. Strikingly, the same covariance structure is present in statistical models of behavioral timing variability [35, 40]. In particular, it matches a Gaussian genrerative model for behavioral interval durations proposed for zebra finch song by Glaze and Troyer [35]. For a set of *P* intervals, this model is parameterized by a vector **w** ∈ ℝ^*P*^ and diagonal latent variable covariance matrices **Ψ** ∈ ℝ^*P* ×*P*^ and **Ω** ∈ ℝ^(*P* −1)×(*P* −1)^. Then, the vector of interval durations in the *µ*^*th*^ trial is generated as

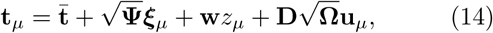

where 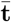 is the average duration and *z*_*µ*_ ∼ 𝒩 (0, 1), ***ξ***_*µ*_ ∼ 𝒩 (**0, I**_*P*_), and **u**_*µ*_ ∼ 𝒩 (**0, I**_*P* −1_) are independent latent factors that are independent and identically distributed across trials. The interval duration covariance matrix in this model is thus

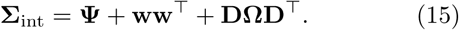

For homogeneously variable intervals, we can therefore associate each component of the covariance matrix in this behavioral model to one of the components of the first-spike interval covariance matrix (13) in our neural model.

The connection between the statistical structures of neural and behavioral variability also leads to a prediction for how variability should scale with behavioral interval duration under a simple timekeeping model. We assume that the basic unit of time is measured via noisy readout of the first-spike times 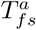 of neurons with covariance (13), and that longer behavioral intervals *T*_*iK*_ are formed by summing the durations of sequences of *K* first-spike intervals:

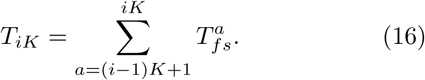

Then, assuming for convenience that *K* evenly divides *N*, the interval-interval covariances are given by sums of the corresponding *K* × *K* submatrices of **Σ**_*fs*_. This summation yields a *N/K ×N/K* interval duration covariance matrix

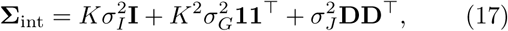

where the scaling of the independent and global components is trivial and that of the jitter component follows from the observation that **DD**^⊤^ is a tridiagonal matrix with sub- and super-diagonal elements equal to −1, first and last diagonal elements equal to +1, and other diagonal elements equal to +2. Therefore, in this simple model for tracking longer intervals of time, the assumption that the first-spike interval covariance matrix has the given structure implies that the interval covariance matrix will have the same structure. Furthermore, the different components of variability have distinct scalings with interval length: the independent component scales linearly, the global component quadratically, and the jitter component is constant. We note that this scaling of local and global components of variability is independent of the details of the single-neuron model, provided that the covariance of the first-spike intervals produced has the form of (13). This variance decomposition and scaling with interval length are illustrated in Figure 3.

### D. Numerical simulations

We compare the theoretical asymptotics we obtained in Sections II A, II B, and II C to the results of numerical experiment. To study the scaling of local variability with noise variance and synaptic strength as shown in Figure 2, we perform 10^4^ realizations of a single-neuron simulation. In these experiments, we fix *τ* = 20 ms, *I*_0_ = − 70 mV, and *V*_*th*_ = − 45 mV while varying *σ* and *I*_*s*_. We integrate the Langevin equation (1) using the Euler-Maruyama method [41] augmented by the reset rule with a timestep of Δ*t* = 10^−3^ ms. The Euler-Maruyama stochastic integration method is an explicit first-order accurate method in the absence of noise, and accurate to order 1*/*2 in the presence of noise. We find empirically that increasing or decreasing the timestep by factors of ten does not influence the qualitative results. For all parameter values tested, we observe good agreement between our asymptotic approximation and the experimental results for the mean first-spike-interval. As shown in Figure 2, for the lowest synaptic strengths and largest noise variances, we observe some discrepancy between asymptotic theory and experiment for the first-spike-interval variance. This is unsurprising, since in that regime *I*_*s*_ +*I*_0_ − *V*_*th*_ is only around five times greater than the standard deviation of the noise, hence higher-order terms in the expansion are non-negligible (see Appendix A 2).

To study the influence of introducing neural fatigue as described in Section II B, we simulate a chain of 80 identical neurons using the methods described above. In these experiments, we fix *σ* = 1 mV and *I*_*s*_ = 45 mV. Over the 10^4^ realizations performed, we draw the parameter *m* from the discrete uniform distribution on {0, …, 249}, with the spiking threshold increment set in terms of the baseline threshold *V*_*th*_ as *δV*_*th*_ = 10^−3^*V*_*th*_. We then define intervals of varying lengths by grouping together uniformly randomly sampled sequences of neurons as described in Section II C. We fit the generative factor model described in Section II C to the intervals generated by our network using the expectation-maximization algorithm described in [35]. In Figure 3b, we plot the square root of the local and global components of variability as a function of interval duration to more clearly illustrate their scaling. We observe good qualitative agreement between the theory and the results of the numerical experiments.

## III. TRIAL-TO-TRIAL TIMING VARIABILITY IN A NOISY HOMOGENEOUS SYNFIRE CHAIN

In Section II, we considered a chain of single neurons with simplified dynamics and coupling for the sake of analytical tractability. In this section, we study variability in a more realistic neural network, a synfire chain [26, 27, 42–44]. A synfire chain is a feed-forward network of multiple pools of neurons, also termed nodes or layers, which are linked by excitatory synaptic connections. We model the neurons in the synfire chain as bursting neurons, and add a set of readout neurons. Bursts have been known to stabilize synfire chains [30], however, we note that the structure of variability does not change if we model neurons that emit a single spike rather than bursts.

We construct a chain of *N* pools, each composed of *M* identical neurons. Every neuron in a given pool *i* is connected to all neurons in the next pool *i* + 1 with equal weights. As before, we approximate the external input to each neuron *I*(*t*) by the sum of an average part *g*(*t*) and a zero-mean Gaussian process 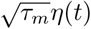 with autocorrelation ⟨*η*(*t*)*η*(*t*′) ⟩ = *σ*_neuron_*δ*(*t* − *t*′). Neurons in the first pool receive an extra input *J* (*t*), which is modeled as a rectangular pulse of height *J*_0_ and width *T*_*p*_. In addition to the per-neuron noise, we include another noise term, which we refer to as “shared” noise. This noise *ξ*(*t*) is generated by a white Gaussian process but is shared across all neurons in a given pool, with mean ⟨*ξ*(*t*) ⟩ = 0 and autocorrelation ⟨*ξ*(*t*)*ξ*(*t*′) ⟩ = *σ*_pool_*δ*(*t t*′). Our motivation for including this additional noise term will become clear later. Then, the sub-threshold dynamics of the membrane potential of neuron *j* in pool *i* are given by

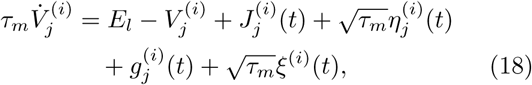

where 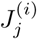 is zero for all *i >* 1. The synaptic input 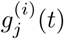 is modeled by the low-pass-filtered spike train [37]

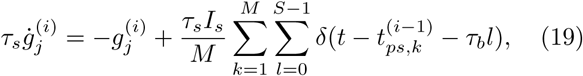

where 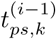 denotes the first-spike time of the *k*^*th*^ neuron of the (*i* − 1)^th^ pool and *I*_*s*_ sets the strength of synaptic coupling. The neurons are modeled to emit a burst of *S* spikes separated by a fixed interval *τ*_*b*_, rather than a single spike. When a neuron’s membrane potential reaches the firing threshold *V*_*th*_, it is then fixed at that threshold until the specified burst duration has elapsed, at which point it is reset to *V*_*r*_, and once again evolves according to the sub-threshold dynamics (18). As in Section II B, we model neural fatigue as a small increase in the membrane potential threshold of all the neurons in the chain after each trial. We also considered a different neural mechanism for fatigue, a simplified model of synaptic depression. However, the structure of the resulting trial-to-trial timing variability is independent of which neural mechanism of fatigue we implement.

For each pool in the chain, we have a single readout neuron, which receives synaptic input from all neurons of that pool along with a white Gaussian noise input with mean zero and autocorrelation *σ*_readout_*δ*(*t* − *t*′) (Figure 1a). The dynamics of membrane potential and synaptic inputs for the readout neurons are similar to that in (18) and (19) with the appropriate inputs.

We study the statistical structure of timing variability in this synfire chain model using numerical simulations. As for the simple model (II D), we integrate the Langevin dynamics (18), (19) using the Euler-Maruyama method, with a timestep of 10^−2^ ms. The parameters values we use are not unique, and are chosen to be in a biologically plausible range. Unless otherwise noted, we simulate a chain of *N* = 81 pools of *M* = 32 neurons each. We set the reset and resting potentials to − 70 mV, the baseline spiking threshold to − 45 mV, and the synaptic strength to *I*_*s*_ = 45 mV. We fix the membrane constant *τ*_*m*_ to 20 ms, the synaptic time constant *τ*_*s*_ to 5 ms, the number of bursts *S* to 4, and the spacing of bursts *τ*_*b*_ to 2 ms. Unless otherwise noted, we let the strengths of the per-neuron, per-pool, and readout noise be 0.5 mV, 1 mV, and 3 mV, respectively. As in Section II D, we fix the spiking threshold increment *δV*_*th*_ to 10^−3^*V*_*th*_, and draw the multiplicative increment factor from the discrete uniform distribution on {0, …, 249} for each trial. Propagation in a synfire chain is not always successful [27, 44]. We consider a trial to be successful if the total number of spikes in the chain is between 4*N* and 4*N* + 0.1 (4*N*), and if all readout neurons fire once. In the experiments on which Figures 4, 5, and 6 were based, two trials out of 1000 were excluded from our simulations that included all sources of noise. In Figure 7c no trials were excluded from the inset.

**FIG. 4.**
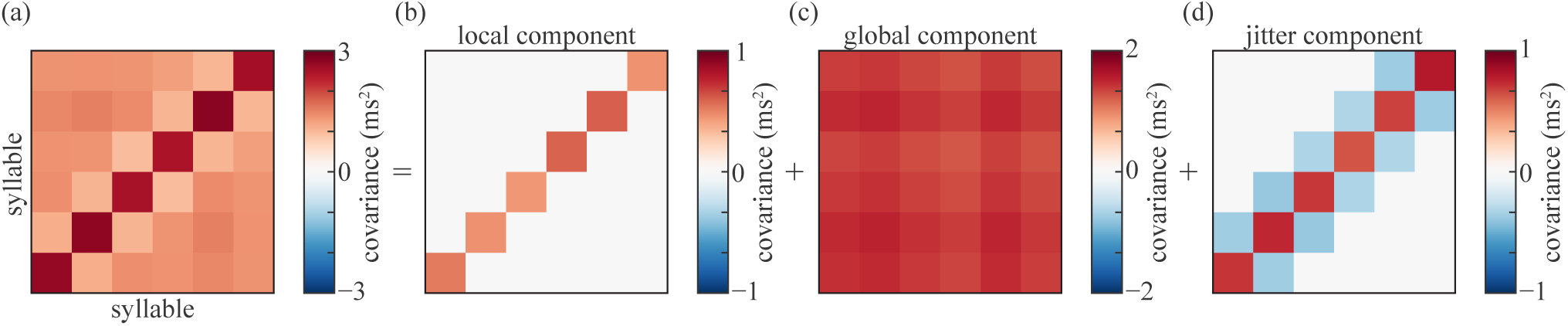
Factor decomposition of the interval duration covariance matrix of the noisy homogeneous synfire chain model. (a) Full model covariance matrix. (b-d) The covariance matrices of the latent factors resulting from applying the analysis method of [35] to the full model shown in (a).

**FIG. 5.**
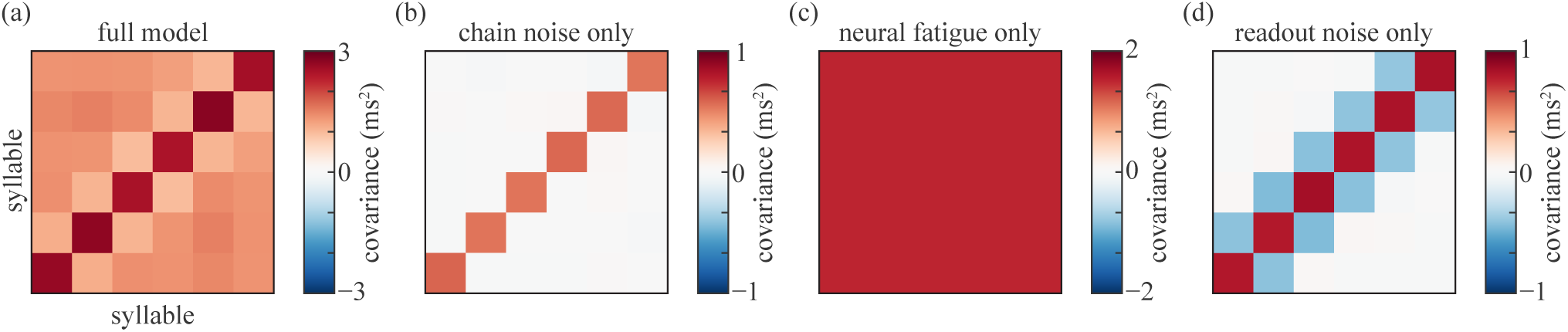
Interval duration covariance matrices of the noisy homogeneous synfire chain model. (a) Interval duration covariance matrix for the full model. (b-d) Interval duration covariance matrices due to chain noise, neural fatigue, and readout noise alone, respectively. In Figure 7 (a,b) we show the contribution of each of the noise sources in (18) to the interval duration covariance matrix separately.

**FIG. 6.**
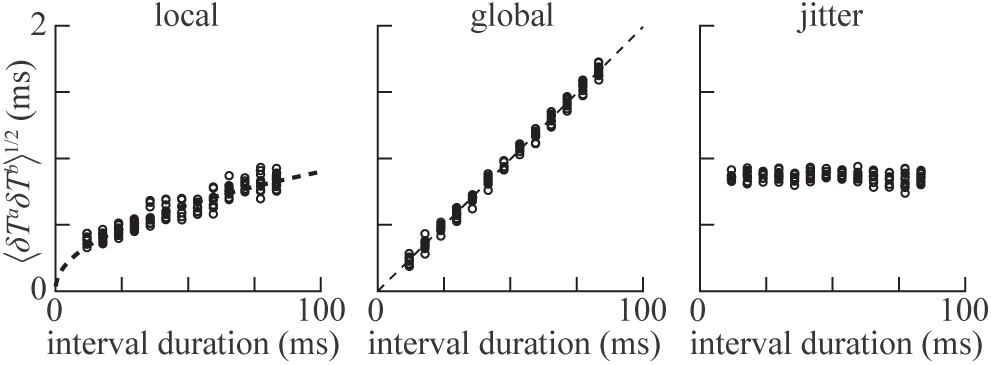
Scaling of local, global, and jitter components of variability with interval duration in the noisy homogeneous synfire chain model with 32 neurons per pool. Circular markers indicate the results of numerical simulations, and dashed lines show power-law fits to the data, with exponents of 0.46 and 1.00 for the local and global components, respectively. No fit is shown for the jitter component, as there does not exist a statistically significant Spearman correlation between it and the interval duration (*ρ* = −0.09, *p* = 0.25).

**FIG. 7.**
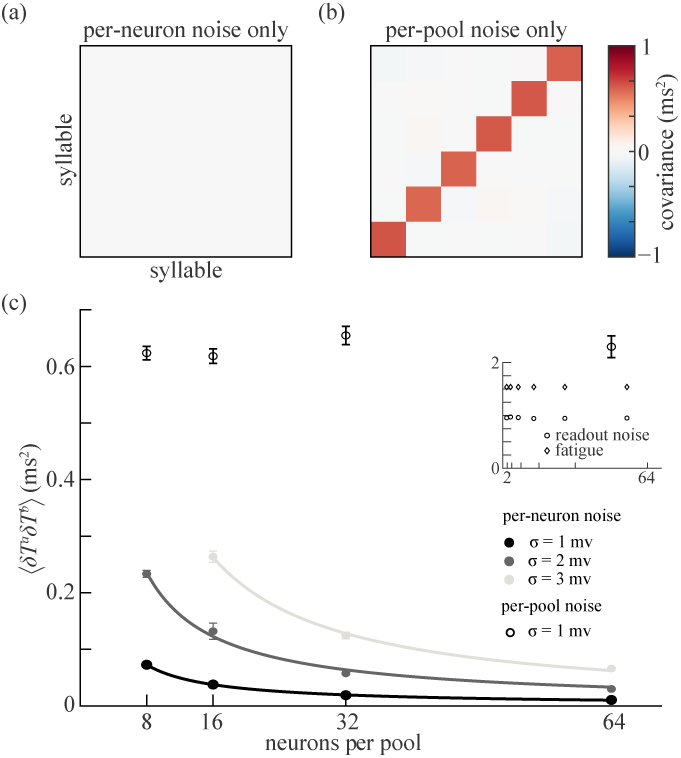
Noisy homogeneous synfire chain model: (a) and (b) show the contribution of each of the noise sources in (18), the per-neuron noise and the per-pool noise which is shared across neurons in the same pool, to the covariance matrix of interval duration.(c) Scaling of interval variability with the number of neurons per pool due to the two noise sources, the per-neuron noise (filled circles) and shared noise among neurons in the same pool (open circles). The filled black, gray, and light gray circles show the interval variability due to per-neuron noise with *σ* = 1 mV, 2 mV, and 3 mV, respectively. Solid lines are power-law fits to the data, with exponents of −0.95 (black line), −0.94 (gray line) and −1.04 (light gray line). Open circles show the same thing but for noise that is shared among neurons in the same pool with *σ* = 1 mV. Error bars show the standard error of the mean. The data point for per neuron noise of *σ* = 3 mV and *M* = 8 was excluded because the chain propagation failed in more than 10% of trials

### A. Relating neuronal mechanisms to different components of trial-to-trial timing variability

In Section II B, we showed that introducing correlated variability to a chain of IF neurons through a simple model of neural fatigue yields a spike interval covariance matrix that is the sum of a local component and a rank-one global component. To test whether this structure is present in the trial-to-trial timing variability of the noisy homogeneous synfire chain model, we define intervals by grouping together sequences of ten neurons, yielding intervals with a mean duration of 59.5 ms and the covariance structure shown in Figure 4 a. Then, we take the full model covariance matrix and use the generative model proposed in [35] (see Section II C) to decompose it into three components, a local component, a global component, and a jitter component, Figure 4. We found that the resulting decomposition explained the covariance matrix well, with a standardized root mean squared residual of 0.0067 [35]. Thus, the statistical structure of trial-to-trial timing variability in the noisy homogeneous synfire chain model is consistent with that of the simple model, with the addition of the component corresponding to readout noise.

Next, we delineate the neural mechanisms behind the components of variability by selectively including different sources of noise. First, we include only the chain noise, which comprises of the shared and the neuron-specific noise terms in the input to a neuron shown in (18). In this case, we recover a diagonal covariance matrix, which we identify as local variability Figure 5b. When we include only neural fatigue, we recover the global component of variability producing a nearly rank-1 covariance matrix, Figure 5c. If we only include noise in the readout neurons, the duration of neighboring intervals are anti-correlated (Figure 5d), corresponding to jitter. This exercise allows us to identify distinct cellular and synaptic mechanisms that explain distinct components of temporal variability: chain noise contributes to the local component, fatigue to the global component, and readout noise to the jitter component.

### B. Scaling of the components of variability with interval duration

In Section II C, we observed that, if one groups multiple neurons together to form intervals, the local component of variability should scale with the square root of the interval length, while the global component should scale linearly with interval length. This scaling is independent of the details of the model, and simply follows from the assumption that the total trial-to-trial variability of the spike interval can be decomposed into a local component and a global component, both of which are uniform in magnitude across neurons. Applying the factor analysis method introduced in [35] to the spike times produced by the noisy homogeneous synfire chain model (Figure 4), we find that the scaling of local and global components of variability with interval length is consistent with this prediction (Figure 6).

### C. Scaling of local variability with pool size

In Figure 7a,b, we show the contribution of each of the noise sources in (18), the per-neuron noise and the per-pool noise which is shared across neurons in the same pool, to the covariance matrix of interval duration. If we vary the number of neurons per pool *M*, we find that the interval duration variance due to per-neuron component of chain noise falls as 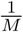 (Figure 7c). Thus, despite the fact that the system is nonlinear, such noise adds quasi-linearly. Therefore, to have a non-negligible local component of variability in the noisy homogeneous synfire chain model, we must assume that neurons belonging to the same pool receive shared noise. In Figure 7c, we see that the interval variability due to this noise mechanism is roughly independent of the number of neurons per pool. Varying the number of neurons per pool did not have an effect on the readout noise or fatigue. This is shown in the inset of Figure 7c.

## IV. THE STRUCTURE OF VARIABILITY IN A BIOLOGICALLY REALISTIC SYNFIRE CHAIN MODEL OF HVC IS CONSISTENT WITH THAT OBSERVED IN ZEBRA FINCH SONG

Zebra finch song is a behavior for which a synfire chain is thought to be involved in the timing of [45]. Zebra finch songs consist of several introductory notes, followed by a few renditions of a motif, sung in a very repetitive manner. Motifs contain 3 to 8 syllables. Syllables range from 50-100 ms and are separated by gaps. The timing of the song is controlled by clock-like bursting in the premotor nucleus HVC, in particular in HVC neurons projecting to Robust Nucleus of the Arcopallium (RA). Many studies suggest that the underlying neural circuit behavior is consistent with a synfire chain model [13, 14, 32, 46, 47]. Further, experimental evidence supports millisecond scale correlations between HVC activity and the song [16, 18, 20]. Thus, we want to test if the trail-to-trail variability in a synfire chain is also consistent with the trial-to-trial variability observed in the song duration. Detailed studies of the trial-to-trial variability in the highly stereotyped zebra finch song were done by Glaze and Troyer [33–35].

We simulate a biologically realistic HVC synfire chain model introduced in [32], which has been shown to agree with measurements of neural variability in HVC. We provide a detailed description of this model in Appendix B. This model consists of a sequentially connected chain of 70 pools, each containing 30 HVC_RA_ neurons, along with a population of 300 HVC_I_ inhibitory interneurons. HVC_RA_ neurons are modeled as two-compartment bursting neurons incorporating dendritic calcium spikes, while HVC_I_ neurons are modeled as single-compartment nonbursting neurons. Inhomogeneity is introduced by randomizing the existence and strength of connections between neurons. Briefly, HVC_RA_ neurons connect to HVC_I_ neurons with probability 0.05, and HVC_I_ neurons connect to HVC_RA_ neurons with probability 0.1. Each HVC_RA_ connects to an HVC_RA_ neuron in the next pool with probability *P* = 0.5 and a connection strength drawn from the uniform distribution on [0, *g*_EEmax_*/*(30*P*)]. Each neuron receives noisy synaptic input in the form of Poisson spike trains which constitutes the chain noise for this model. Synaptic fatigue was modeled by modifying *g*_EEmax_ as (1 − *mδg*)*g*_EEmax_, where *δg* = 10^−3^, with *g*_EEmax_ = 3 mS/cm^2^, and *m* drawn i.i.d. across trials from the uniform distribution on {0, …, 249} as before. All remaining model parameters values are set to those used in Long *et al*. [32]. We read out timing information from the chain using the readout model introduced in Section III. The model was integrated using the Euler-Maruyama method with a timestep of 5 × 10^−3^ ms.

We first observe that the full model covariance matrix of interval duration of this model (Figure 8 a) is similar to that of the homogeneous synfire chain model (Figure 5 a), and has the same structure and magnitude as the song syllable interval duration covariance matrix reported by Glaze and Troyer [35]. Glaze and Troyer [35] showed that the generative model given in (14) is a good description of the statistical structure of the zebra finch song interval duration variability. We previously showed that the structure of variability of the noisy homogeneous synfire chain model is well-described by this generative model (Section III); we find the same to be true for the HVC synfire chain model (Figure 9). The magnitude of variability in the model for 50 ms interval is of order 1 ms (for each component), which is consistent with experimental findings (Figure 10; data from [35]). As in Section III A, we use the results of Section II C to connect behavioral variability to neural mechanisms. Consistent with our results in Section III A, we find that the chain noise contributes to the local component of the song variability, fatigue contributes to the global component, and readout noise contributes to the jitter component (Figure 8).

**FIG. 8.**
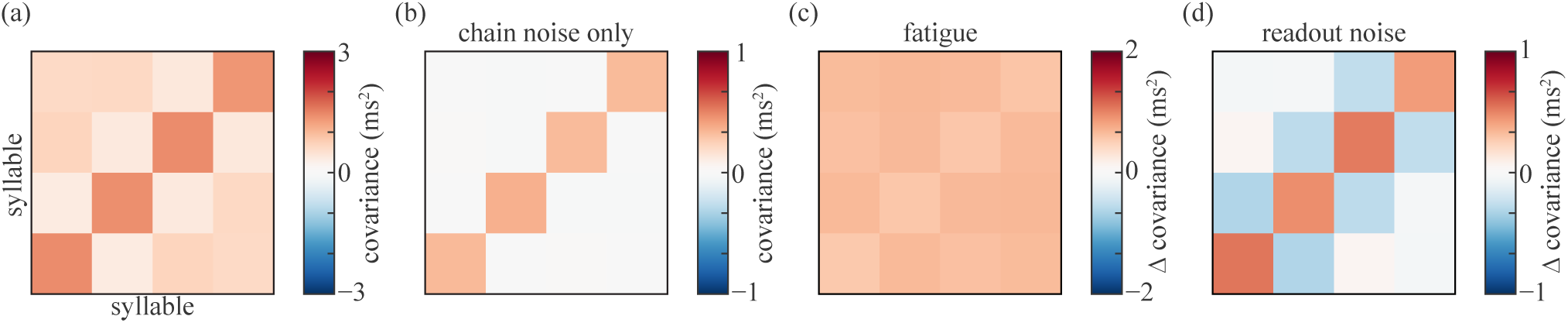
Contributions of different sources of noise in the HVC synfire chain model to the interval duration covariance matrix (model modified from Long *et al*. [32]). (a) Interval duration covariance matrix for the full model. (b) Interval duration covariance matrix with chain noise only. (c) Difference between covariance matrix with chain noise and fatigue and that with chain noise only. (d) Difference between covariance matrix with readout and chain noise and that with chain noise only.

**FIG. 9.**
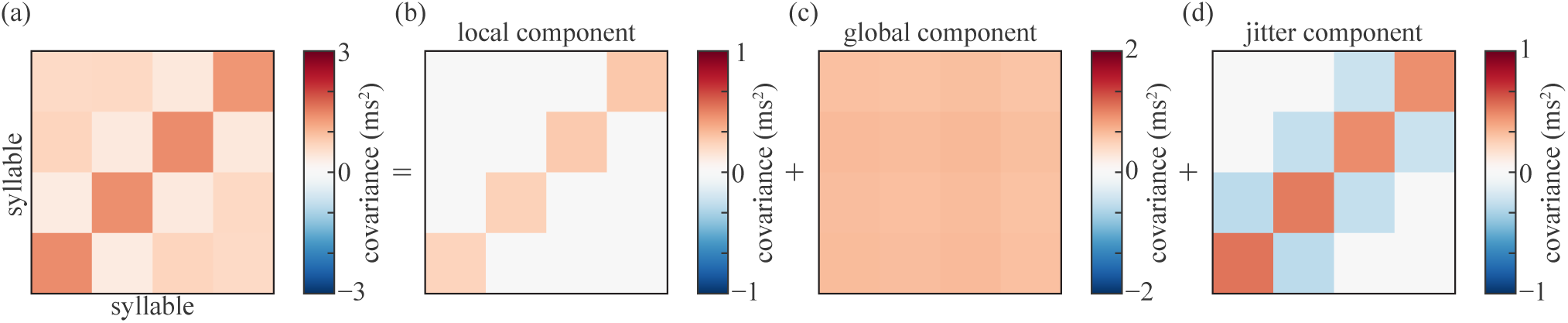
Factor decomposition of the interval duration covariance matrix of the HVC synfire chain model (model modified from Long *et al*. [32]). (a) Full model covariance matrix. (b-d) The covariance matrices of the latent factors resulting from applying the analysis method of [35] to the full model shown in (a).

**FIG. 10.**
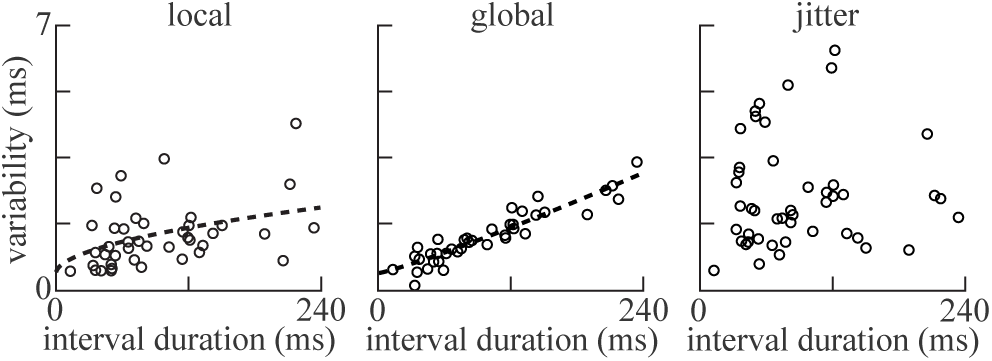
Scaling of the local, global, and jitter components of syllable timing variability with interval duration in zebra finch song (data from Glaze and Troyer [35]). Glaze and Troyer [35] recorded songs of zebra finches, which are composed of a stereotyped sequence of syllables and gaps, a different sequence for each bird. After identifying syllables and gaps, and their durations in each song, they fitted the generative model described in Section II C to this dataset, separately for each bird. Reported data in the figure is extracted from their Figure 3. Each data point represents a syllable-bird pair. As in Figure 6, we fit the relationships between interval duration and the local and global components of variability with power laws, yielding exponents of 0.53 and 1.14, respectively. No fit is shown for the jitter component, as it is not significantly correlated with interval duration.

We also examine how the different components of variability in syllable duration scale with syllable duration. The scalings of the components of variability of syllable duration (Figure 11) agree with the predictions of Section II C, as do previous models (Figures 3b and 6) and the experimental data (Figure 10).

**FIG. 11.**
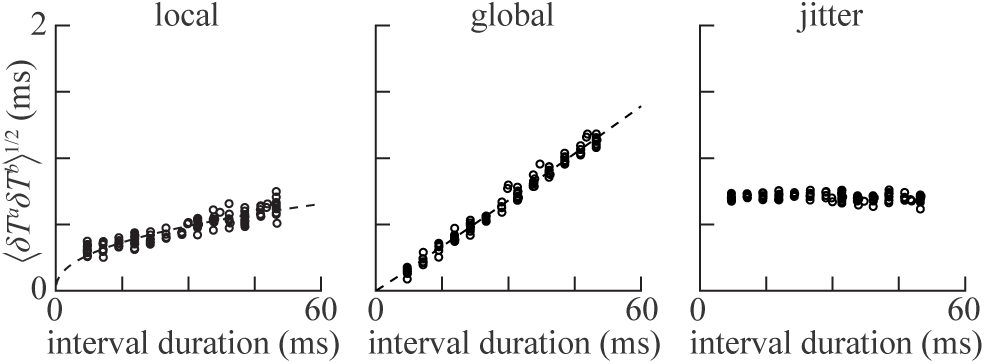
Scaling of the local, global, and jitter components of interval duration variability in the HVC synfire chain model (model modified from Long *et al*. [32]). As in Figure 6, we fit the relationships between interval duration and the local and global components of variability with power laws, yielding exponents of 0.42 and 1.03, respectively. No fit is shown for the jitter component, as it is not significantly correlated with interval duration.

Finally, we note that unlike the noisy homogeneous synfire chain model (Section III), there is no need to add per-pool noise, which is shared across neurons in the same pool. That per-pool noise was necessary in the noisy homogeneous synfire chain model to obtain a non-negligible local component of variability because the variability in input currents to different neurons in a pool would otherwise be uncorrelated (Section III C). In the HVC synfire chain model, sufficient local variability is created by variable synaptic inputs due to uncorrelated presynaptic Poisson spike trains and correlated noisy inputs from inhibitory neurons connecting to multiple neurons in a pool.

## V. DISCUSSION AND CONCLUSION

In this paper, we presented analytical and computational analyses of the trial-to-trial timing variability in synfire chains. We first show how variability scales with input strength and noise level in a simple, analytically tractable chain of single neurons in a low firing rate regime. We also show how trial-to-trial variability scales with interval duration in this simple model. Then, we demonstrated with simulations that our main results carry to noisy homogeneous synfire chains. We found that the statistical structure of timing variability in the chain is well-described by a generative model which consists of local, global and jitter components. Furthermore, we were able to relate each of these components to distinct neural mechanisms in the model. Finally, we show that our main results hold in a biologically realistic synfire chain model of the premotor nucleus HVC in songbirds, and that the structure and magnitude of variability in the model agree with that observed in songbirds song.

Our findings have important implications for the relationship between neural and behavioral variability. Even the most stereotyped of animal behaviors, like the songs of zebra finches, show significant trial-to-trial variability [34]. This variability can be an unavoidable nuisance, or it might be there for an advantageous reason, for example to allow the system to explore more of the behavioral space to help in trial-and-error motor learning [1–3] or help in social contexts [48]. Therefore, understanding the mechanisms which generate and regulate trial-to-trial variability has gained considerable interest [2, 4, 7].

Particularly, it has been argued that some of this variability is rooted in neural activity that controls behavior [2, 4, 7]. Measurements of neural activity show significant trial-to-trial variability [2, 4, 7, 49]. These variations are not independent of behavior; for example, they are known to correlate with behavioral choice in a trial-by-trial basis [50]. Given this background, it is natural to ask whether temporal variability of behavior should reflect the statistics and structure of temporal variability of neural circuits that represent or govern the behavior’s timing in a trial-to-trial basis [9, 12]. Indeed, interrelationships between the timings of neural dynamics and behavior have been observed in various experimental studies [13–20]. For example, Srivastava *et al*. [18] show that in Bengalese finches millisecond-scale changes in the timing of a single spike in a burst in respiratory muscle fibers predicts significant differences in breathing dynamics and millisecond-scale variations in precisely-timed electrical stimulation of respiratory muscles strongly modulate breathing output. This millisecond scale spike timing control of behavior extends to other animals and behaviors [20].

As the temporal variability in zebra finch song and in HVC neurons are both on the millisecond scale [13, 15, 32–35], we speculated that they may be linked. We showed that the temporal variability observed in a biologically realistic model of the zebra finch HVC chain [32] is consistent with the magnitude and structure of the timing variability in the zebra finch song. Thus, our findings provide an example of a detailed match between neural and behavioral variability, and suggest a possible neural account of behavioral variability. A direct experimental test of this suggestion would be to look for correlations in a trial-by-trial basis between millisecond scale temporal variations in the song and spike times of RA projecting HVC neurons.

Potential weaknesses of our neural explanation of the zebra finch song variability are the following. First, while the dominant hypothesis for the song-timing circuit is a synfire chain [32], this question is not fully settled. Further, recent work suggests that the song-timing circuit may not even be fully localized within the HVC but is distributed across multiple areas [51]. If the distributed circuit is a synfire chain, which is consistent with the results of Hamaguchi *et al*. [51], our results still remain valid. Second, in zebra finches, variability of song can be actively regulated through involving lateral magnocellular nucleus of the anterior neopallium (LMAN) [52]. Indeed Ali *et al*. [15] observed that LMAN lesions lead to a reduction in the local component of song timing variability, which was speculated to be mediated by indirect LMAN input to HVC [53]. Another possible source of variability is sensory inference errors [5]. HVC receives feedback auditory input through the nucleus interfacialis of the nidopallium [54], and altered auditory feedback can lead to temporal changes in the song [55]. In the HVC synfire chain model [32], noise was introduced as excitatory and inhibitory independent Poisson spike trains, with no specific reference to where such trains may come from and how could they be regulated. Third, the song production pathway goes from HVC to RA, and then from RA to the tracheosyringeal part of the hypoglossal nerve (nXIIts), which then controls muscular contractions of the syrinx. Neural variability in all these areas as well as variability in muscular contractions contribute to temporal variability of the song. In our model, all of this pathway’s contributions to temporal variability are incorporated into the noise injected to a readout neuron, which in turn contributes mostly to temporal jitter. It is very possible that other components of variability are affected by the downstream activity. Finally, a notion of tempo variation that we did not consider arises from structural changes to the chain, such as homeostatic and synaptic plasticity [15, 46, 47], or experimental perturbations [14]. In birdsong, these mechanisms can lead to tempo changes on the order of tens of milliseconds [14, 15, 46, 47], and, when naturally occurring, require thousands of song repetitions to take effect [15].

**TABLE I.** HVC_RA_ model parameters.

| Name | Description | Value |
| --- | --- | --- |
| $A_s$ | area of somatic compartment | $5000 \mu\text{m}^2$ |
| $A_d$ | area of dendritic compartment | $10000 \mu\text{m}^2$ |
| $C_m$ | membrane capacitance | $1 \mu\text{F}/\text{cm}^2$ |
| $R_c$ | compartment coupling resistance | $55 \text{M}\Omega$ |
| $G_L$ | leak conductance | $0.1 \text{mS}/\text{cm}^2$ |
| $G_{Na}$ | Na<sup>+</sup> conductance | $60 \text{mS}/\text{cm}^2$ |
| $G_{Kdr}$ | delay-rectified K<sup>+</sup> conductance | $8 \text{mS}/\text{cm}^2$ |
| $G_{Ca}$ | high-threshold Ca<sup>++</sup> conductance | $55 \text{mS}/\text{cm}^2$ |
| $G_{CaK}$ | Ca-dependent K<sup>+</sup> conductance | $150 \text{mS}/\text{cm}^2$ |
| $E_L$ | leak reversal potential | $-90 \text{mV}$ |
| $E_{Na}$ | Na<sup>+</sup> reversal potential | $55 \text{mV}$ |
| $E_K$ | K<sup>+</sup> reversal potential | $-90 \text{mV}$ |
| $E_{Ca}$ | Ca<sup>++</sup> reversal potential | $120 \text{mV}$ |
| $E_I$ | inhibitory reversal potential | $-80 \text{mV}$ |

**TABLE II.** HVC_I_ model parameters.

| Name | Description | Value |
| --- | --- | --- |
| $C_m$ | membrane capacitance | $1 \mu\text{F}/\text{cm}^2$ |
| $G_L$ | leak conductance | $0.1 \text{ mS}/\text{cm}^2$ |
| $G_{\text{Na}}$ | Na+ conductance | $100 \text{ mS}/\text{cm}^2$ |
| $G_{\text{Kdr}}$ | delay-rectified K+ conductance | $20 \text{ mS}/\text{cm}^2$ |
| $G_{\text{KHT}}$ | high-threshold K+ conductance | $500 \text{ mS}/\text{cm}^2$ |
| $E_L$ | leak reversal potential | $-65 \text{ mV}$ |
| $E_{\text{Na}}$ | Na+ reversal potential | $55 \text{ mV}$ |
| $E_K$ | K+ reversal potential | $-80 \text{ mV}$ |
| $E_I$ | inhibitory reversal potential | $-75 \text{ mV}$ |

## ACKNOWLEDGMENTS

Part of this work was done when D. Obeid was at the Center for Theoretical Neuroscience at Columbia University. C. Pehlevan acknowledges support from the Intel Corporation. A subset of the computations in this paper were run on the FASRC Cannon cluster supported by the FAS Division of Science Research Computing Group at Harvard University. We thank the anonymous reviewers for a thorough and constructive review.

## Appendix A: Asymptotics for the chain of integrate-and-fire neurons

### 1. Stationary membrane potential distribution in the low-rate limit

Here, we review the approximate firing rate and stationary membrane potential distribution in the low-rate limit *V*_*th*_ − *I*_0_ ≫ *σ* [36]. Inspecting the equation

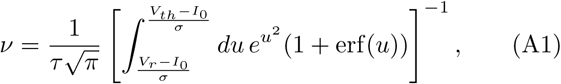

we can see that the integral is dominated by the upper limit due to the exponential. Making the change of variables *u*′ ≡ *u/h, h* ≡ (*V*_*th*_ −*I*_0_)*/σ, v* ≡ (*V*_0_ −*I*_0_)*/*(*V*_*th*_ −*I*_0_), we have

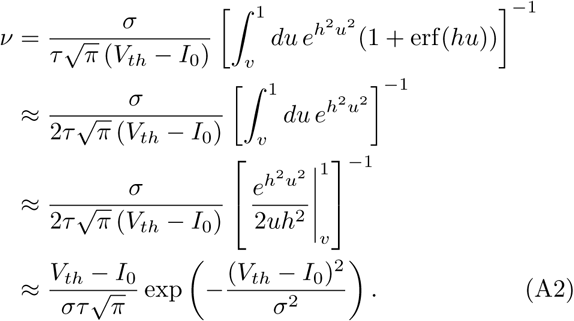

where we integrated by parts in the third line. By a similar argument, we can approximate the stationary distribution of the membrane potential as

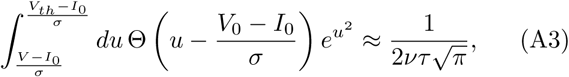

in this limit.

### 2. Moments of the first-spike-interval in the low-rate stationary limit

To derive the moments of the first-spike-interval in the low-rate stationary approximation, we start with the standard results (given as (6) and (7) in the main text) for the mean and variance conditioned on *V*_0_ [36, 37]:

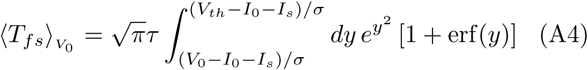

and

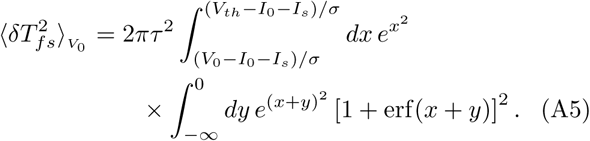

Considering the mean first-spike-interval, we use the integral representation of the error function as

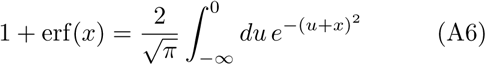

to write

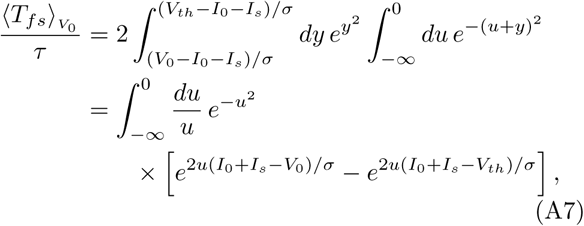

where we note the cancellation in the bracketed integrand that ensures that it does not diverge as *u* → 0^−^. We can then easily compute the expectation over the approximate distribution (5) of *V*_0_ to obtain

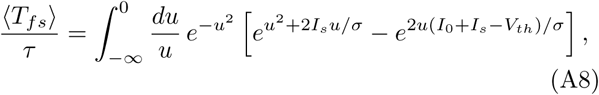

which, though it does not have a simple closed-form solution, is the integral of a bounded entire function that decays exponentially fast at infinity (provided that *I*_*s*_ *>* 0), and is therefore well-behaved.

To obtain a similar integral expression for the variance of the first-spike-interval, we recall the law of total variance

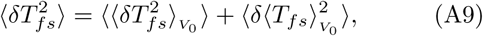

where the outer angle brackets denote averaging over the distribution of *V*_0_, and follow the same procedure that we used to derive ⟨*T*_*fs*_⟩ to obtain

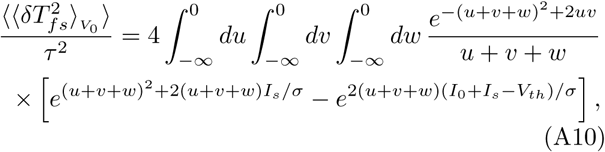

and

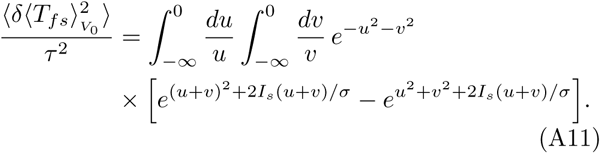

With these integral expressions in hand, we can now derive asymptotic expansions for the moments. For brevity, we define the dimensionless scalars *α* ≡ (*I*_0_ + *I*_*s*_ − *V*_*th*_)*/σ* and *β* ≡ (*V*_*th*_ − *I*_0_)*/σ*; we will work in the limit of low baseline firing rates *β* ≫ 1 and large synaptic inputs *α* ≫ 1. Rescaling *u* by 2*α* in (A8), we can write the mean first-spike-interval as

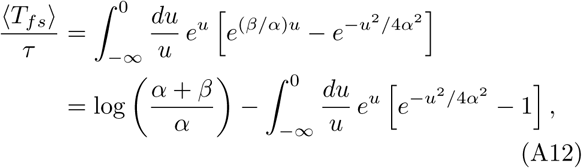

where we have split the integral into two pieces by adding and subtracting one from the integrand and evaluated the first of the remaining integrals. Expanding the remaining integrand other than the overall exponential weight *e*^*u*^ as a power series and integrating term-by-term using the relationship of the integrand to the gamma function [56], we obtain the divergent asymptotic series

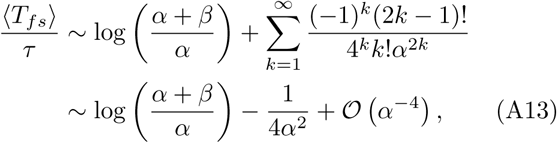

which yields the lowest-order approximation given in the main text.

We now consider 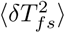. Converting the integral over the negative octant in (A10) to an integral over the positive octant, making the change of variables *x* ≡ *u, y* ≡ *v* + *w, z* ≡ *w*, and parameterizing the domain of integration such that we integrate first over *z* ∈ [0, *y*], we have

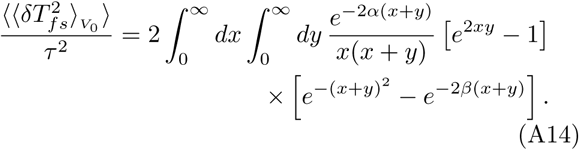

Then, adding the expression for 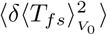 given in (A11) to the above expression for 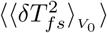 as prescribed by the law of total variance (A9), we have

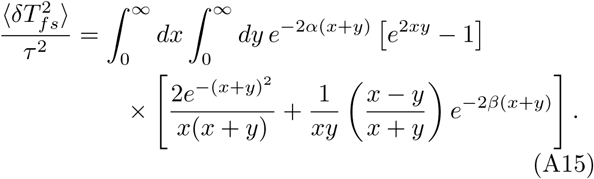

As it is anti-symmetric about the line *y* = *x*, the second term in the bracketed integrand will vanish under integration over the positive quadrant, leaving

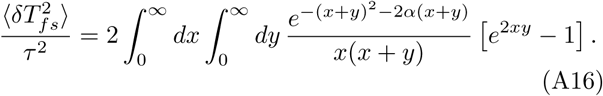

Rescaling *x* and *y* by 2*α* and making the change of variables *u* ≡ *x* + *y, v* ≡ *x*, we have

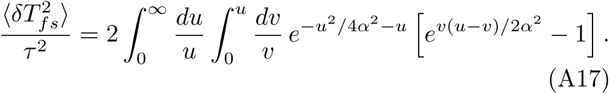

Expanding the bracketed portion of the integrand as a power series and observing that

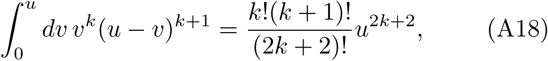

we have, integrating over *u* term-by-term,

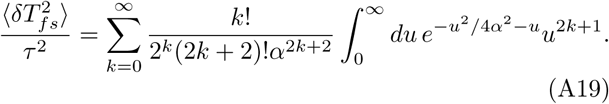

To allow us to apply standard asymptotic results to the remaining integral, we note that it is related to Tricomi’s confluent hypergeometric function *U* (*a, b, z*) as *α*^2*k*+2^(2*k* + 1)! *U* (*k* + 1, 1*/*2, *α*^2^) [56], hence, shifting indices for convenience, we can write

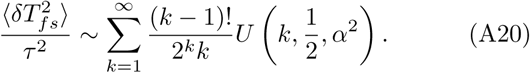

Using the standard result that

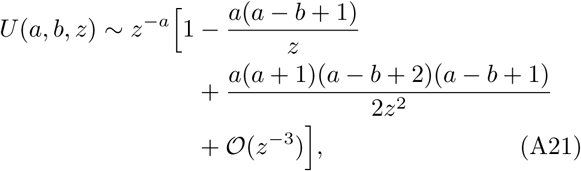

for |*z*| ≫ 1 [56], we have

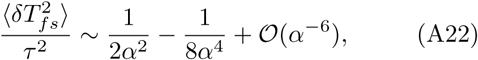

which yields the lowest-order approximation given in the main text.

To obtain the asymptotic approximations for the timing variability in the simple model for neural fatigue given in the main text (10, 11, 12), we start from the asymptotic expansions without fatigue (8, 9), and apply the laws of total expectation and total variance given the assumed distribution of the parameter *m*. We then expand the resulting expressions about the baseline spiking threshold *V*_*th*_ to lowest order in *δV*_*th*_*/*(*I*_*s*_ + *I*_0_ − *V*_*th*_), assuming that *m*_max_ *δV*_*th*_ ≪ *I*_*s*_ + *I*_0_ − *V*_*th*_, yielding the asymptotic approximations (10) and (11).

### 3. Moments of the first-spike-interval in a delta-function approximation

In the previous appendix and in the main text, we considered the approximation of the distribution of initial membrane potentials by the stationary Gaussian limit (5). In this appendix, we consider a delta-function approximation *P* (*V*_0_) ≈ *δ*(*V*_0_ − ⟨*V*_0_⟩). This approximation maps directly to the standard treatment of leaky IF neurons with the appropriate replacement of *V*_*r*_ by *I*_0_. Here, we review the derivation of the corresponding asymptotic results [37, 38]. In the limit *V*_*th*_ − *I*_0_ ≫ *σ* of low firing rates, we have *(V*_0_) = *I*_0_, hence we fix *V*_0_ = *I*_0_ in this approximation. Considering the mean first-spike-interval, we again start from the standard expression (6) with *V*_0_ set to *I*_0_, and rescale *σy 1*→ *y*, yielding

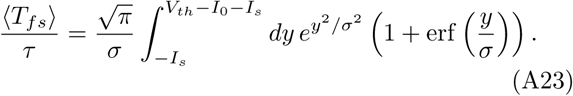

In the limit *I*_*s*_ + *I*_0_ ≫ *V*_*th*_ *σ* of large synaptic inputs, the quantity *y* in the above integrand is always negative, and we have *y/σ* ≪ −1. Using the asymptotic expansion of the error function for *x* ≪ −1 [56],

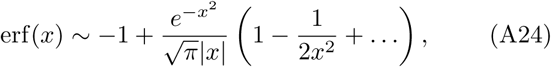

we obtain

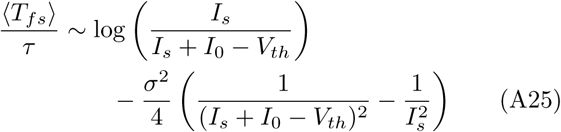

to lowest order. Similarly, for the variance of the first-spike-interval, we start with the standard expression (7) with *V*_0_ = *I*_0_. Again rescaling the variables of integration by *σ* and using the asymptotic form of the error function, we obtain the lowest-order approximation

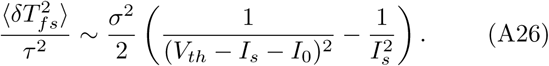

Comparing these expressions to the corresponding results (8, 9) in the approximation of the initial membrane potential distribution by the stationary Gaussian distribution (5), we observe that they are identical up to the presence of the 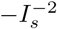 terms in the lowest-order approximations. The presence of these terms in the delta-function approximation means that the variability decreases more rapidly with increasing synaptic strength and increases less rapidly with increasing noise variance *σ*^2^ than in the Gaussian approximation.

## Appendix B: Details of the HVC synfire chain model

In this appendix, we provide a detailed description of the HVC synfire chain model from Long *et al*. [32]. This model consists of a chain of 70 sequentially-connected pools of 30 HVC_RA_ neurons, along with a population of 300 HVC_I_ inhibitory interneurons. A given HVC_RA_ neuron connects to an HVC_RA_ neuron in the next pool with probability *P* and an excitatory synaptic conductance drawn from the uniform distribution on [0, *g*_EEmax_*/*(30*P*)] mS cm^-2^, where *g*_EEmax_ is a dimensionless parameter. An HVC_RA_ neuron connects to an HVC_I_ neuron with probability 0.05 and excitatory synaptic conductance drawn uniformly from [0, 0.5] mS cm^-2^. Finally, an HVC_I_ neuron connects to an HVC_RA_ neuron with probability 0.1 and an inhibitory synaptic conductance drawn uniformly from [0, 0.2] mS cm^-2^. Long *et al*. [32] chose these parameter values such that successful spike propagation was possible for many values of *g*_EEmax_.

### 1. HVC_RA_ dynamics

In the Long *et al*. [32] model, HVC_RA_ neurons are modeled as two-compartment bursting neurons. The somatic compartment contains leak, Na+, and delay-rectified K+ conductances, while the dendritic compartment contains leak, high-threshold Ca++, and calcium-activated K+ conductances. This model can generate dendritic calcium spikes, which evoke stereotyped bursts of sodium spikes in the soma. The membrane potentials *V*_s_ and *V*_d_ of the somatic and dendritic compartments obey the dynamics

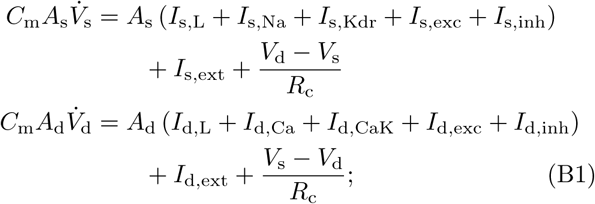

we enumerate the definitions and values of all parameters in Table I. The dynamics of the injected currents *I*_s,ext_ and *I*_d,ext_ are freely chosen, while the remaining currents are given as

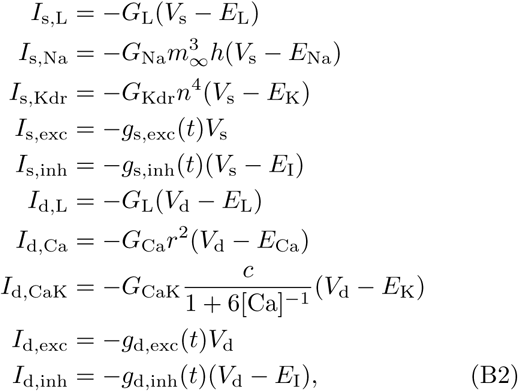

where *g*_s,exc_, *g*_s,inh_, *g*_d,exc_, and *g*_d,inh_ are the total synaptic conductances of the soma and dendrite. The gating variable *m*(*t*) = *m*_∞_(*V*_s_) is an instantaneous function of *V*_s_, while *h, n, r, c* all evolve according to the dynamics

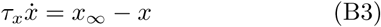

for *x* ∈ {*h, n, r, c*}, where the activation functions are given as

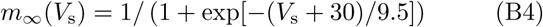

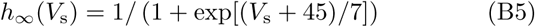

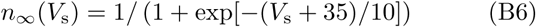

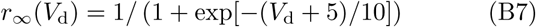

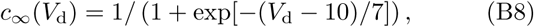

and the time constants are given as

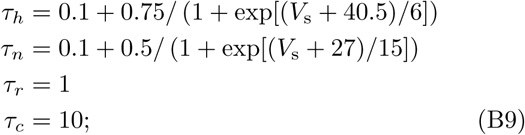

the units of all constants are implied. Finally, the calcium concentration [Ca] evolves as

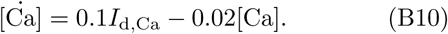

Synaptic conductances follow “kick-and-decay” kinetics: *g* ↦ *g* + *G* when a spike arrives at a synapse with conductance 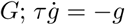 between spikes, for *g* ∈ {*g*_exc_, *g*_inh_ }. The synaptic time constants *τ*_exc_ and *τ*_inh_ are both fixed to 5 ms.

### 2. HVC_I_ dynamics

In the Long *et al*. [32] model, HVC_I_ neurons are modeled as single-compartment neurons containing leak, Na+, delay-rectified K+, and high-threshold K+ conductances. The membrane potential *V* obeys the dynamics

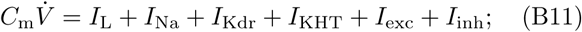

the values of all parameters are given in Table II. The currents are given as

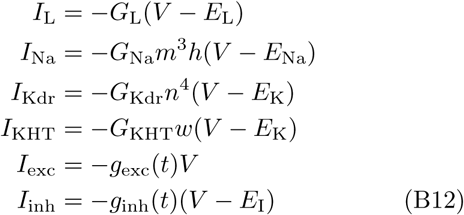

for total excitatory and inhibitory synaptic conductances *g*_exc_ and *g*_inh_. The gating variables *m, h*, and *n* evolve according to the dynamics

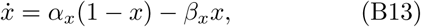

for *x* ∈ {*m, h, n*}, where

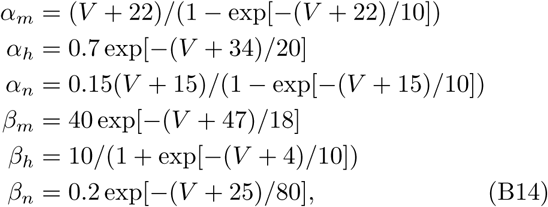

with implied units throughout. The gating variable obeys

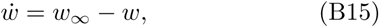

where

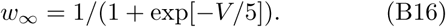

The excitatory and inhibitory conductances obey the same dynamics as for HVC_RA_ neurons, except for the fact that the excitatory time constant *τ*_exc_ is set to 2 ms.

### 3. Noise spike trains

The Long *et al*. [32] model introduces noise into the neurons via independent Poisson spike trains. Each HVC_RA_ neuron receives excitatory and inhibitory spike trains at both compartments, each generated from a homogeneous Poisson process with a rate of 100 Hz. The conductances of each spike are drawn independently in time from a uniform distribution on [0, 0.035] mS*/*cm^2^ for the somatic compartment and [0, 0.045] mS*/*cm^2^ for the dendritic compartment. Each HVC_I_ neuron also receives excitatory and inhibitory noise spike trains, generated from 250 Hz Poisson processes with conductances drawn uniformly from [0, 0.45] mS*/*cm^2^. With this noise model, the RMS fluctuation in the membrane voltage of each compartment of each HVC_RA_ neuron is about 3 mV, and the HVC_I_ neurons spike spontaneously at about 10 Hz.

## Notes

### Competing Interest Statement

The authors have declared no competing interest.

